# Protein engineering of a genetically encoded biosensor for wastewater detection of profen NSAIDs

**DOI:** 10.64898/2026.01.14.699353

**Authors:** Zoë Davis, Hao Tian, Ian Wheeldon, Sean R. Cutler, Timothy A. Whitehead

**Affiliations:** Department of Chemical and Biological Engineering, University of Colorado Boulder; Boulder, 80305, USA; Institute of Integrative Genome Biology, University of California, Riverside; Riverside, CA, 92521, USA; Department of Botany and Plant Sciences, University of California, Riverside; Riverside, CA, 92521, USA; Department of Chemical and Environmental Engineering, University of California, Riverside; Riverside, CA, 92521, USA; Center for Industrial Biotechnology, University of California, Riverside; Riverside, CA, 92521, USA

**Keywords:** Biosensor, non-steroidal anti inflammatory drugs, wastewater monitoring, ketoprofen, protein engineering, deep mutational scanning, protein engineering

## Abstract

Non-steroidal anti-inflammatory drugs (NSAIDs) are pervasive environmental contaminants due to their frequent and widespread use, multiple paths of release into surface and ground water supply, diversity of the chemical class, and toxicity to aquatic and other non-target species. In particular, the 2-arylpropionic acid (“profen”) class of NSAIDs poses significant risks to aquatic ecosystems due to incomplete removal during wastewater treatment. Current monitoring precludes high frequency testing at point sources. Here we present the engineering and application of a genetically encodable, protein-based biosensor for the detection of the NSAIDs ketoprofen and pranoprofen in wastewater effluent. We repurposed the plant hormone receptor PYR1 to bind selectively to profens using computational protein design, deep mutational scanning, and yeast 2 hybrid and yeast surface display screening. The resulting sensor, PYR^NSAID^, has a nanomolar limit of detection for ketoprofen and panoprofen, and µM sensitivity to the NSAIDs ibuprofen, fenoprofen, tolmetin and diclofenac. We also demonstrated dose responsive-activity of our sensor in simulated wastewater matrices containing the common wastewater contaminants sulfamethoxazole, caffeine, acetaminophen, and 2,4, dichlorophenol using a split Nanoluc luminescence assay. PYR^NSAID^ is the first step towards a scalable, cost-effective alternative for real-time monitoring of pharmaceutical pollution.

## Introduction

Non-steroidal anti-inflammatory drugs (NSAIDs) are some of the most commonly prescribed and over the counter drugs used, with thousands of tons per year consumed worldwide^1^. Point sources for NSAID pollution include hospitals, pharmaceutical industrial pollution, sewage from incomplete human metabolism, and landfills or septic tanks. Their frequency of use, paired with prescription across multiple industries, contributes to a difficulty of NSAID clearance from wastewater^2,3^.

Profens, or 2-arylpropionic class of NSAIDs, include the common NSAIDs ibuprofen, ketoprofen/dexketoprofen, fenoprofen, and pranoprofen. Ibuprofen is a well known human medicine. Ketoprofen is commonly distributed to livestock and approved for use in the US as well as many European and Asian countries. Multiple studies have established µg/L concentrations of the common profens in wastewater^4,5^, surface water^6,7^, groundwater^4^, and soil^8^ worldwide. NSAIDs released into the environment pose the greatest risk to aquatic life. Chronic exposure to profens have been found to decrease reproduction, disrupt development, and damage endothelial lining of crustaceans, aquatic microbes, fish, and algae at concentrations in the high nM to low µM range^9,10,11^.

Methods of wastewater screening for NSAIDs vary depending on the treatment plant. A typical protocol for chronic exposure testing involves shipping of water samples to off site facilities and exposing model aquatic organisms for up to a week^12^. In the case of quantitative monitoring, a combination of high performance liquid chromatography and mass spectrometry or diode array detection^13^ is employed off site. These methods are not always reliable in complicated matrices^14^, and requiring transportation of samples also decreases the likelihood that wastewater treatment plants will test at multiple steps in the treatment process, or that they will monitor effluent not planned for drinking use. Ideally, wastewater would be cleared of NSAIDs *before* release into aquatic ecosystems. This would prevent chronic exposure of vulnerable organisms and breakdown of NSAIDs into new toxic products with hard to predict downstream effects.

There has been recent work developing sensors for NSAIDs in wastewater effluent using electrodes ^15,16,17^, ionophores^18^, nanomaterials^19,20^, and mammalian cell based assays to detect overall COX activity^21,22^. While promising, there are some limitations to these methods; electrodes are susceptible to fouling and require frequent calibration that would decrease the potential of sensor adoption for use at the treatment source. In addition, cell based activity assays are ill-suited for the complicated and toxic matrices characteristic of wastewater effluent. Genetically encodable biosensors have been developed for PFAS^23^ and estrogens^24^, and a genetically encodable NSAID sensor could potentially overcome some of these limitations to sensing profens in wastewater effluent. However, to our knowledge, no genetically encodable biosensor for profens has been developed.

PYR1 is a plant hormone receptor that undergoes a conformational change after binding to abscisic acid, in turn enabling it to bind to the phosphatase HAB1^25^. This chemically induced dimerization (CID) mechanism has been engineered for use as a biosensor to detect a wide range of drug-like small molecules, including organophosphates^26^, agrochemicals^27^, herbicides^28^, illicit drugs^29^, terpenes^30^, and steroids and other natural metabolites^31^. This PYR1-HAB CID mechanism can be exploited for a range of screening and sensing modalities^32^, including split luciferase, phosphatase inhibition, and yeast transcriptional assays^26^. The plasticity of the receptor binding pocket and the potential for multimodal signalling outputs enable a wide range of applications in medicine, environmental sensing, and drug screening using both *in vitro* and *in vivo* systems.

In this study, we engineer a nanomolar-responsive PYR1 biosensor–PYR^NSAID^–for the NSAIDs ketoprofen and pranoprofen. PYR^NSAID^ was developed using yeast 2 hybrid (Y2H) selections, yeast surface display (YSD) screening, computational design, and deep mutational scanning. PYR^NSAID^ binds to the propionic acid NSAIDs dexketoprofen, ketoprofen, pranoprofen, and acetic acid NSAID diclofenac with nanomolar sensitivity. PYR^NSAID^ also exhibits micromolar sensitivity for other propionic acid NSAIDs like ibuprofen. Importantly, PYR^NSAID^ shows minimal cross-reactivity with other common wastewater contaminants like caffeine or acetaminophen. We demonstrate profen sensing using a yeast-based assay as well as in vitro using a split luciferase format. PYR^NSAID^ shows promise as a point-of-treatment sensor with environmentally relevant limits of detection for a range of NSAIDs commonly found in wastewater.

## Materials and Methods

### Chemicals

All chemicals used in yeast surface display experiments and the split luciferase assay were ordered from Sigma-Aldrich, including pranoprofen (Cat #SML1423), ketoprofen (#K1751), dexketoprofen (#PHR2626), ibuprofen (#PHR1004), diclofenac (#PHR1144), tolmetin (#1670502), acetaminophen (#PHR1005), caffeine (#PHR1009), sulfamethoxazole (#PHR1126), and 2,4, dichlorophenol (#D70406).

### Plasmid construction

Full starting plasmid sequences can be found in the supporting information. Gene fragment and primer sequences can be found in the supporting information. All variants were cloned into plasmids via Golden Gate assembly^33^. All plasmids were sequenced using Oxford Nanopore (Plasmidsaurus). Plasmid pJS723^26^ was used for ΔN-HAB1^T+^ expression. All yeast genetic libraries were cloned into the yeast surface display plasmid pND003^34^. PYR-LgBit and HAB-SmBit proteins for the split luciferase assay were expressed in plasmids pSDS003^29^ and pSDS036^29^, respectively.

### Protein Purification and Preparation

ΔN-HAB1^T+^ protein was purified, chemically biotinylated, and stored in 100 M saturated ammonium sulfate using the HAB1 purification protocol previously described in Steiner, et al.^34^. For purification of SmBit-HAB1^T+^-MBP and LgBit-PYR variants, the same expression and purification protocol was used^34^, minus the chemical biotinylation step.

### PYR1 Library Design

A computationally inspired library was created from previously described yeast 2 hybrid sensor hits for ketoprofen and pranoprofen^31^. Ketoprofen contains a ketone—similar to the native ligand abscisic acid—and pranoprofen possesses a carboxylic acid, which we hypothesized would hydrogen bond with a bound water bridging PYR1 and HAB1. To determine likely conformers of ketoprofen and pranoprofen, we performed temperature replica exchange MD simulations as previously described^29^. Temperatures ranged from 300 to 450K with replica exchange every 100 ps and run for a total of 100ns. The ligands were parameterized using OpenFF Sage with the TIP3P water model. The source code used is freely available (https://github.com/ajfriedman22/SM_ConfGen). Conformers for each profen were used to replace abscisic acid in the PYR1 binding pocket, and we docked only the *S*-enantiomer of both ketoprofen and pranoprofen. We followed this with a Rosetta repacking and filtering step^29^. Rosetta output was manually curated to revert residues identified as important for PYR-HAB binding and PYR1 yeast display^34^. We also added amino acids of a similar class if multiples from that class were present in designs; i.e., adding all uncharged and small hydrophobic, or large and branched amino acids. The final library included reversion mutations back to the PYR1 parental sequence for most positions; a mutation table can be found in **Table S1**.

### Construction of PYR1 mutational libraries

Combinatorial oligonucleotides were designed using custom Python scripts to include the necessary overhangs and add mutations to the thermally stabilized abscisic acid-binding HOT5 background^35^. These were ordered as a combined oligo pool from Agilent Technologies (oligonucleotide sequences are shown in Supporting_Data1.xlsx). Three different PCRs were performed from the pooled oligos, corresponding to three separate gene-encoding cassettes for PYR. The mutated PYR library was cloned into yeast display plasmid pND003^35^ by Golden Gate assembly^33^. Cassettes for the Golden Gate assembly included the three oligo pool cassettes covering the mutated region and two gene fragment HOT5 cassettes flanking either side of the mutated region. We transformed the library into XL1 Blue electrocompetent cells (Agilent Technologies, Cat No. 2002249) following the manufacturer’s protocol and recovered the transformation on a large square bioassay plate (Thermofisher, Cat No. 240835). Transformation efficiency was determined using serial dilution (**Table S2**). Cell biomass on the large bioassay plate was scraped and plasmid midiprepped using a ZymoPure^TM^ plasmid midiprep kit (Zymo Research, Cat No. D4200). This library was transformed into chemically competent *S. cerevisiae* EBY100^37^ and stored as 1mL, OD_600_ = 1.0 yeast storage buffer (20% w/v glycerol, 200mM NaCl, 20mM HEPES pH 7.5) stocks and kept at −80 °C according to Medina-Cucurella et al^38^.

The site saturation mutagenesis (SSM) library for PYR^prano^ was performed by nicking mutagenesis^39^, using oligonucleotide primers designed by software from Suni mutagenesis^40^. Positions 59, 81, 83, 92, 94, 108, 110, 117, 120, 122, 141, 159, 160, 163, 164, and 167 were mutated to every other amino acid plus stop codon using a NNK degenerate codon. The mutagenized library was transformed into *E. coli* Mach1 chemically competent cells, plasmid extracted and purified, transformed into *S. cerevisiae* EBY100, and stored as noted above.

### Yeast Surface Display Screening

Our yeast induction protocol closely followed the methods outlined in Steiner, et al. 2020^34^ with one change of using 10% SDCAA, 90% SGCAA as the induction media (for 1 L: 2 g dextrose, 18g galactose, 6.7 g Difco yeast nitrogen base, 5 g Bacto casamino acids, 5.4 g Na_2_HPO_4_, 7.44 g NaH_2_PO_4_). Cells were grown in SDCAA at 30 °C to an OD_600_ of 2.0, then centrifuged and resuspended in induction media to grow overnight at 20 °C. In preparation for yeast surface display binding reactions, ammonium sulfate precipitated ΔN-HAB1^T+^ protein was spun down at 17,000g for 10 minutes, resuspended in ice cold CBSF++ (147mM NaCL, 20mM sodium citrate, 4.5mM KCl, 0.1 (w/v)% bovine serum albumin, pH 8.0 with 1mM DTT and 1mM TCEP added the day of experiments) and desalted into the same CBSF++ buffer using Zeba desalting columns (Thermofisher, Cat No. A57759). 100mM ligand stock solutions were made fresh each day of experiments with the following solvents: ketoprofen, pranoprofen, and dexketoprofen were dissolved in DMSO; ibuprofen and fenoprofen were dissolved in ethanol; tolmetin stock solutions were made with nanopure water. Yeast surface display experiments were performed on a Sony SH800S cell sorter (Sony Biotechnology), closely following the protocol in Leonard et al.^29^

### Yeast Display Sorting

For all sorting experiments, growth and induction conditions were identical to those described above. Representative cytograms and gates for this library can be found in **Fig S1.** For screening of the PYR1 computationally designed library, three sorts were performed; these sorting gates and concentrations can be found in **Fig S2**. Ligand concentrations, top binding % sorted, and number of cells collected were as follows. Sort 1: 100 M pranoprofen/ketoprofen, highest 4% of binders for both ligands. Sorted populations for each ligand were screened separately after the first sort. Sort 2: 100 M pranoprofen/ketoprofen, top 8% of pranoprofen binders and top 9% of ketoprofen binders. Sort 3: 10 M pranoprofen/ketoprofen, top 8% of pranoprofen binders and top 23% of ketoprofen binders. For all sorts of the PYR1 computationally designed library, 500nM ΔN-HAB1^T+^ was used. To sort the site saturation mutagenesis library, we used 300nM ΔN-HAB1^T+^, 33.3 M pranoprofen, and 100 M ketoprofen. We also sorted the top 3% of binders for 10 M abscisic acid and an identical gate that collected ∼0.3% of top constitutive binders in the SSM library, combining these two groups for subsequent next generation sequencing analysis. Corresponding cytograms and sorting gates for the site saturated mutagenesis library can be found in **Fig S3**. Once sorted, cells were grown at 30 °C for 36 hours in SDCAA supplemented with 1x penicillin-streptomycin solution (Thermofisher, Cat No. 15140122) and stored as 50 v/v% glycerol stocks in yeast storage buffer at −80 °C.

### Next generation sequencing and sequencing analysis

Sorted populations were grown in SDCAA at 30°C for 36 hours and DNA was extracted using freeze/thaw cycles and zymolase according to the protocol in Medina-Cucurella & Whitehead^38^, with a few changes as noted. PCR reactions were run on a gel to check length, gel extracted with an NEB Monarch DNA Gel Extraction Kit, washed with 500 µL DNA wash buffer, and eluted in 30 µL nuclease free water. Plasmid DNA purified with exonuclease I and lambda nuclease was cleaned using an NEB DNA & PCR cleanup kit (New England Biolabs, Cat No. T1135) according to the manufacturer’s instructions. This purified DNA was amplified with primers flanking the mutagenized region of PYR1 and containing adapter sequences for Illumina sequencing. PCR was performed with 12.5 µL Q5® High-Fidelity 2X Master Mix following the manufacturer’s protocol. The resulting DNA pools (one for each ligand, as well as a negative binding and reference pool) were submitted for Illumina paired end short read sequencing. The resulting reads were merged with FLASh^40^. Using a custom Python script, this data was trimmed and cleaned to only include reads that were the right length and had the correct start and end sequence. For the initial library, this script calculated the highest frequency designs; the top three mutants for each chemical were tested for dose responsive activity via yeast surface display (**Fig S2**). To analyze the site saturation mutagenesis library, we added another step in the NGS data processing code to only include reads containing a single mutation from the parent sequence in the 16 binding site residues we targeted. These NGS reads were used to determine the actual coverage of our site saturated mutagenesis library (**Table S3**), as well as the enrichment of each mutation for all experimental groups (ketoprofen, pranoprofen, no ligand) normalized to the reference (**Fig 3A,B; Fig S6**). Heat maps of enrichment ratios were created by calculating the log_2_(frequency of mutation in experimental group/frequency of mutation in reference) at all mutated positions for Keto^+^, Prano^+^, and negative/ABA^+^ sorts. Mutations with zero read counts were given a pseudocount of 1 so as to avoid a calculated ERROR. Any mutation identities with a read count less than 8 were left out of heat maps and marked as having ‘not enough data’ (**Fig 3A,B; Fig S6**).

### Luciferase Plate Assays

In preparation for split luciferase assay experiments, SmBiT-HAB and LgBit-PYR were desalted and experiments were performed following the protocol in Leonard et al.^29^ with the following changes: ligand concentrations for full titrations ranged from 100nM to 1mM, and % v/v of cosolvent was 2% in each well. Ketoprofen, caffeine, and 2,4, dichlorophenol were dissolved in DMSO; acetaminophen and sulfamethoxazole were dissolved in ethanol. LgBiT-PYR^NSAID^:SmBiT-HAB1^T+^-MBP ratio was optimized for a final assay setup with 5nM LgBiT-PYR^NSAID^ and 50nM SmBiT-HAB1^T+^-MBP in a total well volume of 200 uL.

## Results

### Identification of initial PYR1 biosensors using Y2H and YSD screening

Two of the most common NSAIDs by volume are ketoprofen and ibuprofen^42^ which share a 2-aryl propionic acid common to all profens (**Fig 1A**). Although all profens have this structural similarity, there are differences in chirality and relative hydrophobicity of the aryl substituent (**Fig 1A**). Our goal was to design a sensor capable of binding multiple profens at environmentally relevant concentrations. Based on NSAID aquatic toxicity studies^10,11,41^ as well as ketoprofen environmental monitoring data^6^, we aimed for a sub-1 µM (∼250 g/L) limit of detection for both ketoprofen and pranoprofen.

**Figure 1.**
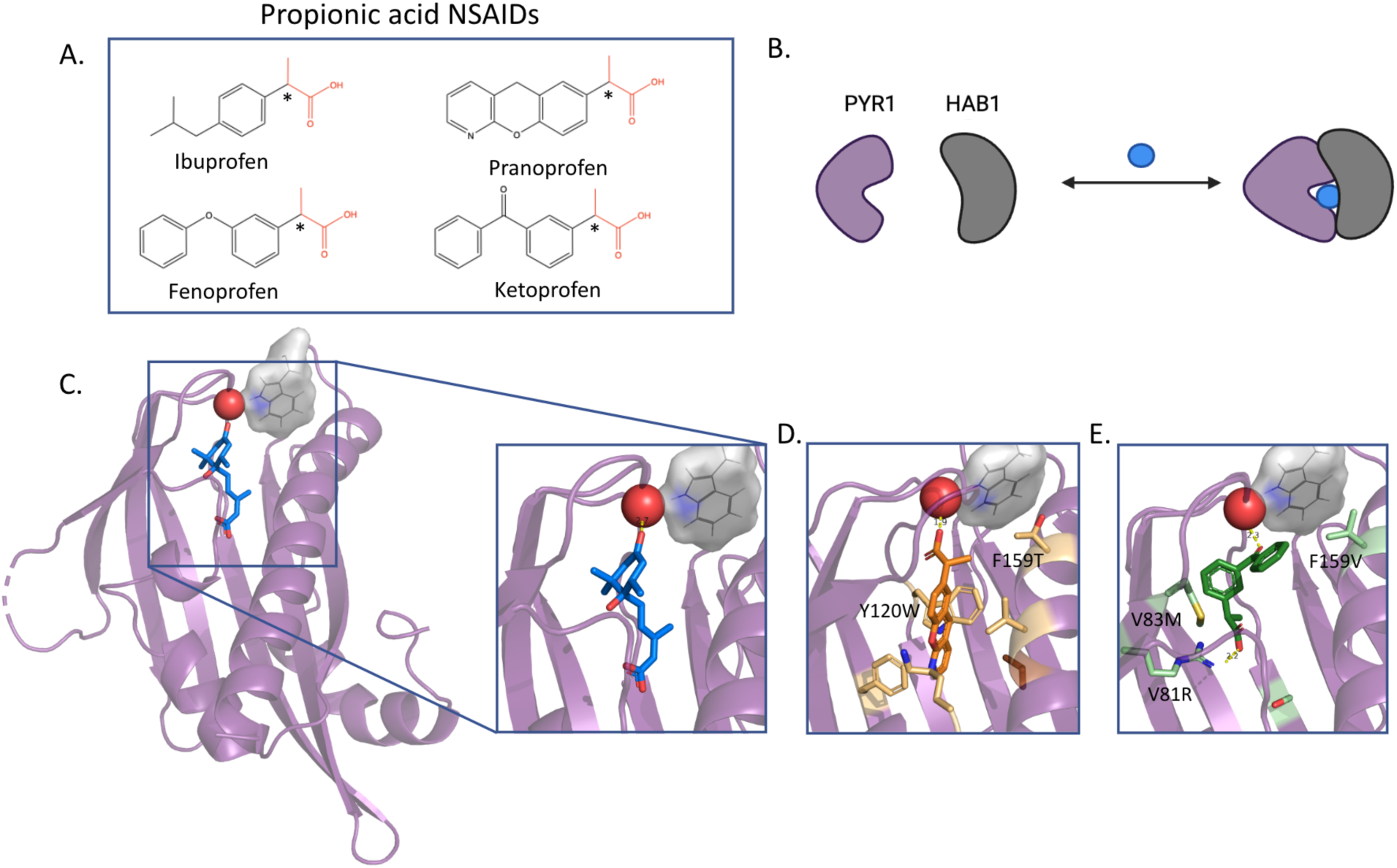
| Computational design of a propionic acid NSAID library using initial PYR1-HAB1 yeast 2 hybrid sensor hits for ketoprofen and pranoprofen binders. **A.** R groups of common propionic class NSAIDs (profens). The asterisk denotes a chiral center; all profens listed are given as racemic mixtures of the two enantiomers. **B.** Cartoon of the plant hormone receptor PYR1 (purple) and its chemically induced dimerization to HAB1 (gray) by a ligand (blue ball). **C.** Experimentally determined structure of the abscisic acid (blue sticks)-PYR1 (purple cartoon)-HAB1 (gray surface and sticks) ternary complex (PDB ID: 3QN1). Only Trp385 of HAB1 is shown. The red sphere at the top of the binding pocket is a bound water critical for the gate-latch-lock chemical induced dimerization mechanism for PYR1-HAB1. Modeled NSAID ligand fits in the PYR binding pocket are shown for **D.** pranoprofen-specific (prano5) and **E.** ketoprofen-specific (keto3) Y2H hits. Profens are shown as **D.** orange (panoprofen) and **E.** forest green (ketoprofen) sticks. Mutations for the hits are shown as wheat and mint sticks for the pranoprofen and ketoprofen sensors, respectively.

The plant hormone receptor PYR1 and phosphatase HAB1 form a dimer upon binding with the native ligand abscisic acid (**Fig 1B**). Previously, a computational PYR1 mutational library was developed for a variety of chemicals^26^ and screened using Y2H to identify ten PYR1-based sensors for ketoprofen (seven) and pranoprofen (three) (**Table S4**)^31^. These sensors were reported to bind near the limit of detection for Y2H (100 M for ketoprofen, 10 M for pranoprofen). The pranoprofen sensors all contain a F159V/T mutation along with 1-2 additional, diverse mutations in the receptor binding pocket. In contrast, the three ketoprofen mutations contained shared mutations at V81R, V83L/M, and F159I/V. These initial Y2H hits were used as starting points for a new library of potential profen sensors.

A bound water connecting abscisic acid, PYR1, and HAB1 is essential to the energetic strength of the ternary complex^41^ (**Fig 1C**). To determine plausible binding modes for the Y2H sensors, we used a previously described ligand docking method^29^ to (i.) generate likely conformers of ketoprofen and pranoprofen using temperature replica exchange molecular dynamics; and (ii.) align a hydrogen bond acceptor on the ligand to the ketone of the native ligand abscisic acid (**Fig 1C, D, E**). In the case of pranoprofen, multiple alignments in the binding pocket were possible due to mutually incompatible mutations in the initial yeast two hybrid hits (**Table S4**). Some mutations increased the space and polarity along the sides of the pocket (F108Q/N), accommodating hydrogen bonding of the internal oxygen in pranoprofen with the bound water molecule. Other mutations included packing along the sides of the binding pocket and increased space and hydrophobicity near the bottom (Y120W, K59L), leading us to believe that the carboxylic acid in pranoprofen was participating in hydrogen bonding with the bound water and the ligand was fitting lengthwise in the active site (**Fig 1D**). In this case, the shared mutation F159V/T could be rationalized as relieving unfavorable interactions between the hydrophobic aromatic and the negatively charged carboxylate. For ketoprofen, we hypothesized that its ketone was a hydrogen bond acceptor to the key bound water molecule (**Fig 1E**). This was based on yeast two hybrid hits all containing a V81R mutation that would allow a salt bridge with the carboxylic acid on ketoprofen. Since our goal was to create a sensor that bound to both NSAIDs, we submitted the central oxygen alignments and the carboxylic acid alignments for both ketoprofen and pranoprofen (**Fig 1D, E, Fig S4 A, B**) to a Rosetta design and filtering step^29^. Additionally, we used rational design to add amino acids from a similar class, different charges and polarities in polar and charged areas of the active site, or to accommodate fit for conflicting alignments (e.g. including all small hydrophobic residues at site 59, several branched hydrophobic residues at 83, multiple polar uncharged branched amino acids at position 117, and two large aromatic residues at 120) (**Table S1**). Finally, for most positions we included the parental PYR1 residue in our library. In total, our 1.4 million member library contains mutations at 15 different positions (**Table S1**).

### Screening of computationally inspired library identifies profen-sensitive sensors

Using yeast surface display, we tested whether our computationally inspired library would contain profen sensors (**Fig 2A**). We assembled our library using Golden Gate assembly^33^, transformed into the yeast display yeast strain EBY100^37^, and prepared stocks for sorting. In total, we obtained an estimated 30.3% coverage of the library members (410,000 estimated unique variants; **Table S2**). This library was sorted for three rounds of fluorescence activated cell sorting^34^ with a biotinylated, thermally stabilized ΔN-HAB1 protein (HAB1^T+^)^26^ and either ketoprofen (100 M sorts 1-2; 10 M sort 3) or pranoprofen (100 M sorts 1-2; 10 M sort 3) (**Figure S2-3**). Binding was measured by secondary labeling with a streptavidin-phycoerythrin fluorescence conjugate. We noted an increase of constitutive binding (binding in the absence of ligand) as the sorts progressed. Still, ligand-dependent binding was clearly observed in sort 3. After the third sort, ketoprofen and pranoprofen populations were collected and sequenced by short read Illumina NGS. For each population, three variants comprised the majority of the sequencing reads (**Table S5**). We ordered synthetic genes encoding these variants and tested them for profen-specific binding using yeast surface display. All variants showed dose-dependent responses (**Figure S5**). As expected from the results from the third sort, most variants had high constitutive activity. Still, we identified a PYR1 variant from the pranoprofen sorts (PYR^prano^) with 7 mutations (K59N, V83L, E94M, S122Q, F159T, A160V, V164E) that had low constitutive binding, high binding signal with 100 M pranoprofen (**Fig 2B**), and statistically significant binding signal in the presence of 100 M ketoprofen (**Fig 2C**). At this concentration, PYR1^prano^ showed higher binding signal than the top sensors identified initially by Y2H screening (**Fig 2B, C**). In order to optimize NSAID binding affinity, PYR^prano^ became the parent sequence for a subsequent deep mutational scanning optimization of the sensor pocket.

**Figure 2.**
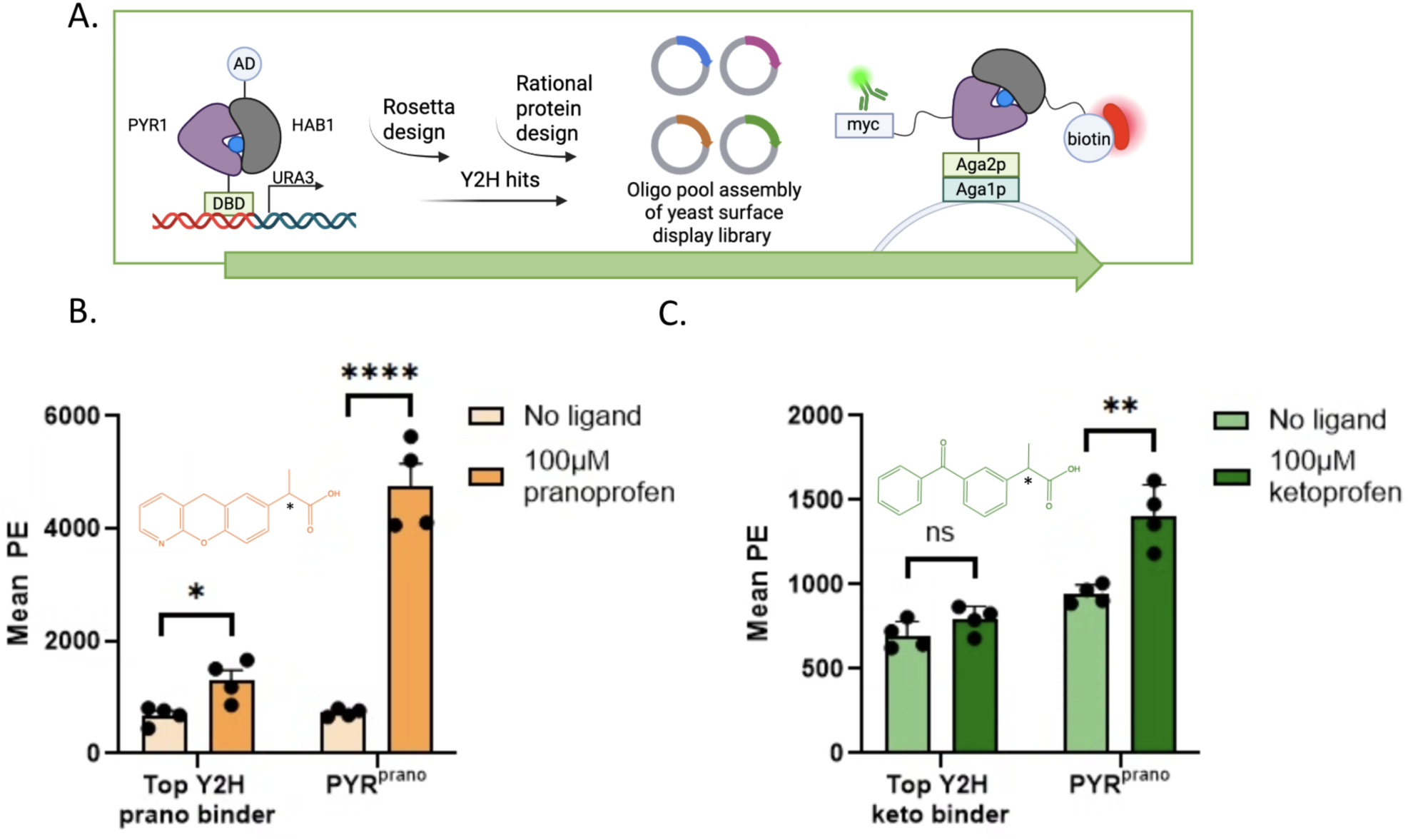
|A lead sensor binding both ketoprofen and pranoprofen identified from yeast surface display screening of a computationally-inspired library. **A.** Flow chart of identification of a lead sensor hit through yeast 2 hybrid (Y2H) and yeast surface display (YSD) screening of computationally inspired PYR1 mutational libraries. Y2H hits were used as input for a new PYR1 mutational library, which was screened by YSD. **B.** Yeast surface display binding to ketoprofen and pranoprofen for the top PYR1 hits after Y2H and YSD screening. Mean PE binding signal was measured using a biotinylated 200 nM ΔN-HAB1^T+^, followed by secondary labeling with streptavidin-phycoerythrin (SAPE). For all data, n = 4. ns, not significant; *, p-value < 0.05; ** p-value < 0.01; *** p-value <0.001.

### Optimization of initial ketoprofen and pranoprofen sensor by deep mutational scanning

To improve PYR^PRANO^ for binding to pranoprofen and ketoprofen, we performed deep mutational scanning using yeast surface display coupled to fluorescence activated cell sorting. We screened a focused single site saturation mutagenesis (SSM) library at 16 positions in the PYR1 binding pocket (**Table S3, 6**). This SSM library was sorted under four conditions: (i.) a reference sort on the displaying population only; (ii.) a sort collecting the top 2% of the binding population at 100 µM ketoprofen and 300 nM HAB1^T+^ (Keto^+^); (iii.) a sort collecting the top 5% of the binding population at 33.3 µM pranoprofen and 300 nM HAB1^T+^ (Prano^+^); and (iv.) a sort collecting the top 1% of the binding population containing 300 nM HAB1^T+^ and no other ligand (Constitutive^+^) (**Fig S3**). After sorting, plasmids from the four yeast populations were extracted and amplicons were prepared comprising the portion of the PYR1 gene corresponding to the locations of the mutations. These amplicons were sequenced using Illumina paired end short read sequencing. Sequencing read counts for each population were converted to an enrichment ratio (e.r.; a log_2_ transformed frequency change) by comparing the binding populations (Keto^+^, Prano^+^, Constitutive^+^) to the reference population. Heatmaps of the enrichment ratios for the Keto^+^ and Prano^+^ sorts are shown in **Figure 3A** and **3B**, respectively. Mutations which were enriched were largely shared between sorts (**Figure 3C**). In total, 18 mutations were identified that had an e.r. above zero for both the Keto^+^ and Prano^+^ sorts.

**Figure 3.**
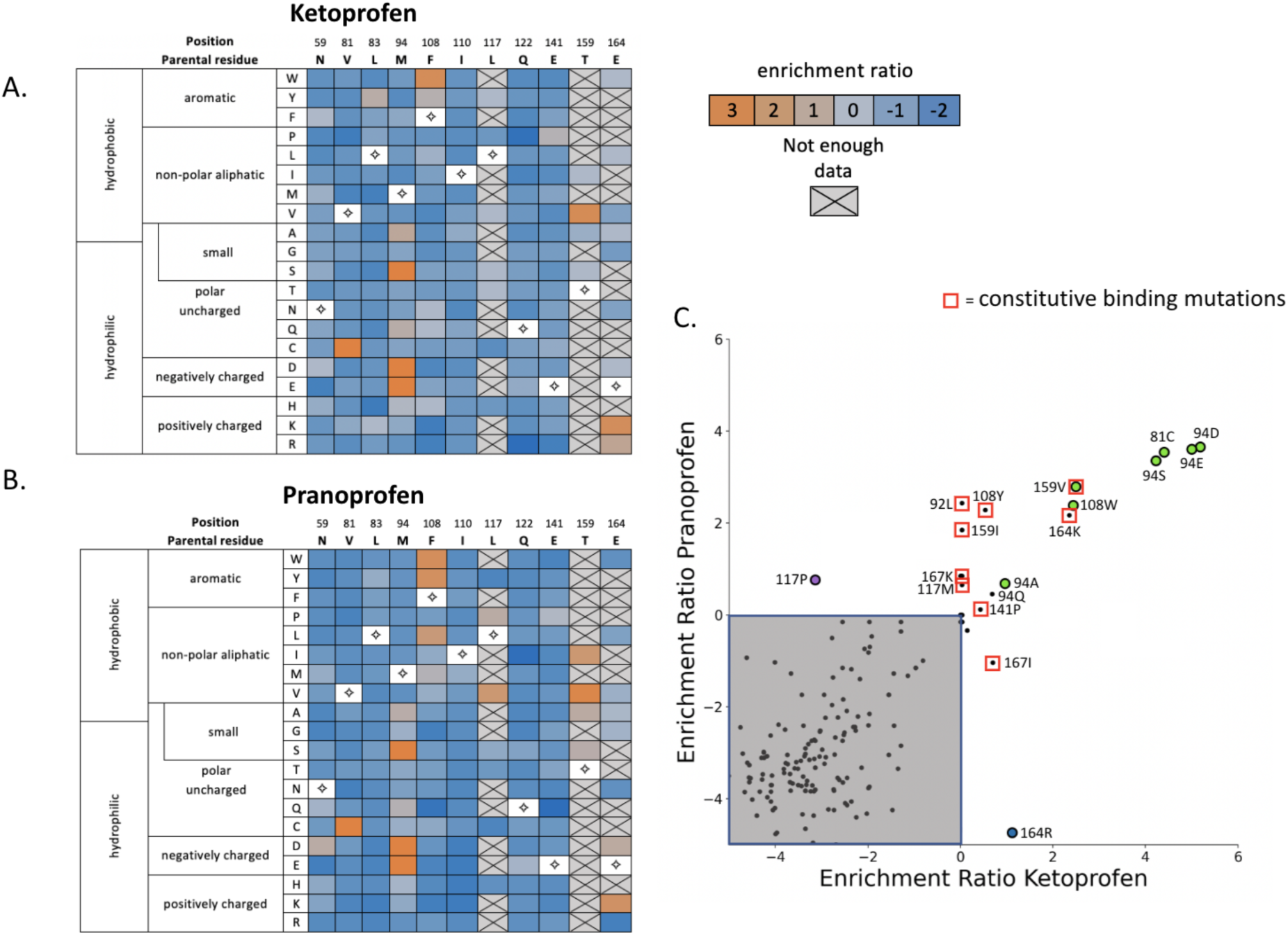
| Deep mutational scanning of PYR1^prano^ against ketoprofen and pranoprofen. **A., B.** Heatmap showing enrichment ratios at 11 positions in the PYR1 binding pocket. Yeast display sorts were performed using 300 nM ΔN-HAB1^T+^ and (**A.**) 100 M ketoprofen or (**B.**)33.3 M pranoprofen. Enrichment ratios were determined using a reference sort of no ligand and screening using gates for protein expression only. White boxes with diamonds represent the parental sequence for the PYR^PRANO^ sensor. **C.** Enrichment ratios for pranoprofen plotted against ketoprofen. The gray box demarcates enrichment ratios below zero for both ligands. The open red boxes show mutations that are constitutive (binding in the absence of ligand), as determined by deep mutational scanning. Mutations marked with colored circles were tested in combination to identify PYR1^NSAID^. Green, blue and purple circles represent pan, ketoprofen, and pranoprofen binding enriched mutations, respectively.

The mutational profile indicated areas where the local sequence was optimized, and positions where the design residue was sub-optimal. Of the residues shared between PYR1 and PYR^prano^, V81C (e.r. 4.4 for Keto^+^, 3.5 for Prano^+^) and aromatic mutations at position 108 (F108WY) showed strong enrichment. Of the seven mutations in PYR^prano^ compared with PYR1, three (N59, L83, Q122) had no mutation with a positive enrichment ratio in both sorts. Another position (V160) did not have sufficient coverage to determine its optimality. For position 159, a mutation of threonine to its isostere valine is enriched in both Keto^+^ and Prano^+^ populations, but it also is enriched in the Constitutive^+^ population. Notably, in the isolation of PYR^prano^, the other candidates with high constitutive background all contained the V159 mutation. The remaining two positions (94, 164) showed that charge modulating mutations were strongly enriched. The highest enrichment ratios for the Keto^+^ and Prano^+^ sorts occurred for a reversion of M94E (**Fig 3C**), with nearly the same enrichment for M94D. The non-conservative mutations M94A/S were also enriched. For position 164, a charge reversal E164R was enriched in the Keto^+^ but not the Prano^+^ population. Finally, a non-conservative mutation (L117P) was depleted in the Keto^+^ population and moderately enriched for the Prano^+^ population.

We sought to combine mutations identified from the deep mutational scan to improve PYR^prano^ for profen recognition. We kept any hits with an experimental group (Keto^+^, Prano^+^) e.r. > 0.5 and an e.r. < 0.5 in the no ligand group; 7 mutations across 4 different positions fit these criteria (**Fig 3C**). We also tested the mutations L117P and E164R that were indicated as mutations specific for pranoprofen and ketoprofen binding, respectively. We performed this optimization in two steps. In the first step, we used yeast display to screen 12 of the 16 possible combinations of 94ADES, 108FW, and 159TV, leaving out the 4 variant combinations mutated only at site 94.

We screened for constitutive binding and binding in the presence of 1 mM ketoprofen or 33.3 M pranoprofen (**Figure S7**). 159V was associated with higher constitutive binding compared with 159T, consistent with the screening results at this position observed for the isolation of PYR^PRANO^ (**Figure S5**). Designs including serine at position 94 hit had slightly lower constitutive activity and higher ketoprofen binding signal than the reversion mutation to glutamate (**Fig S7**). Additionally, the mutation F108W increased binding to ketoprofen and pranoprofen (**Fig S7**). In the second step, we constructed and tested a second round of mutational combinations in the background of PYR^PRANO/M94S/F108W^(**Fig S8**). From this round, we additionally identified V81C and V164R as viable mutations for our final sensor.

### Validation of a nanomolar responsive profen sensor in yeast

We next assessed PYR1^NSAID^ (PYR1^HOT5^ K59N, V81C, V83L, E94S, F108W, S122Q, F159T, A160V, E164R) for binding affinity and specificity using yeast display. PYR^NSAID^ contains a total of nine mutations from its parental background - five mutations from PYR^PRANO^ and four from SSM optimization. In this assay, PYR^NSAID^ exhibited a limit of detection (LOD) of 330nM and 770nM for ketoprofen and pranoprofen, respectively (**Figure 4B**; LODs were calculated using the 3σ method, which is equivalent to the drug concentration that yields a signal equal to 3 times the standard deviation of the blank after subtraction). Notably, PYR^NSAID^ has a much improved responsiveness in this yeast assay compared with PYR^PRANO^ and the top hits from the Y2H screen (**Figure 4B**). Ketoprofen exists as a racemic mixture; dexketoprofen, the (S)-enantiomer of ketoprofen, has a LOD of 50 nM using the same yeast assay, suggesting preferential recognition of this enantiomer (**Figure 4C)**. To determine if PYR^NSAID^ had potential as a more general profen and acetic acid NSAID sensor, we also tested PYR^NSAID^ binding to ibuprofen, fenoprofen, and the acetic acid NSAIDs diclofenac and tolmetin (**Figure 4C-D**). While binding profiles were significantly weaker than for ketoprofen and pranoprofen, PYR^NSAID^ had a LOD of 1 µM for diclofenac (**Figure 4C**) and statistically significant binding activity at 33.3 µM ibuprofen and 10 µM fenoprofen and tolmetin (**Figure 4D**).

**Figure 4.**
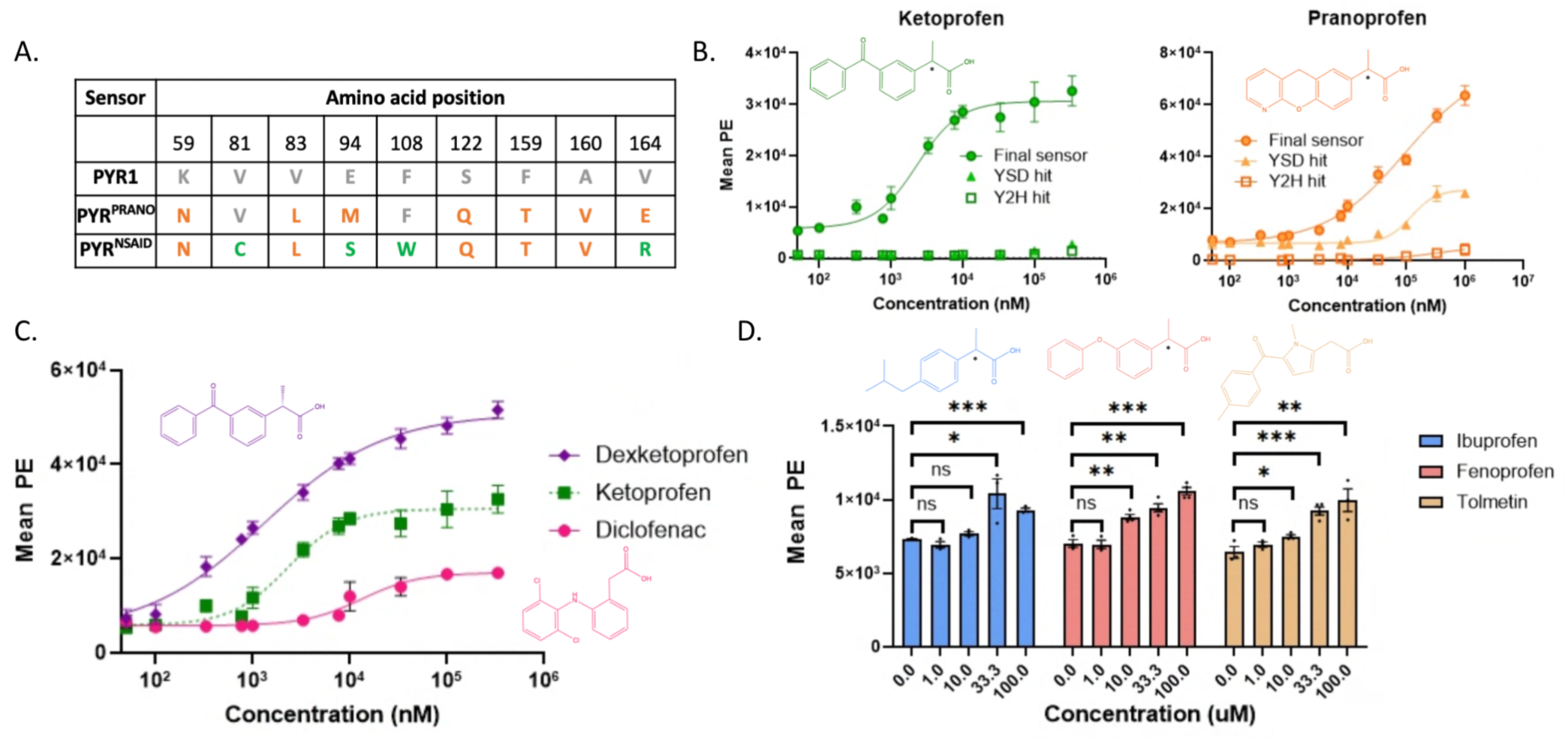
| A nanomolar responsive sensor for profen NSAIDs. **A.** Residues for selected positions in the PYR1 binding pocket for different sensors. PYR1^NSAID^ is the engineered final sensor. **B.** Yeast surface display titrations for ketoprofen (left panel) and pranoprofen (right panel) for PYR1^NSAID^ (closed green circles), PYR1^PRANO^ (closed triangles), and the top hit for Y2H (open squares). **C.** Yeast surface display titrations for other propionic acid NSAID dexketoprofen (purple diamonds) and acetic acid NSAID diclofenac (pink circles). All PYR proteins were tested in the thermostable HOT5 background. **D.** Mean PE vs concentration for PYR1^NSAID^ for the indicated NSAIDs. Mean BE binding signal was measured using a biotinylated 200 nM ΔN-HAB1^T+^, followed by secondary labeling with streptavidin-phycoerythrin (SAPE). For all data, n=4. Error bars represent 1 s.e.m. and in some cases may be smaller than the symbol. ns, not significant; *, p-value < 0.05; ** p-value < 0.01; *** p-value <0.001.

### An *in vitro* assay for assessing profen water contamination

We next tested whether PYR^NSAID^ would selectively bind to NSAIDs, even in the presence of other chemicals commonly found in wastewater matrices. As representative effluent water contaminants we chose caffeine, the antibiotic sulfamethoxazole, the triclosan degradation product 2,4, dichlorophenol, and acetaminophen. These chemicals were chosen for their widespread use and persistence in the water column^45^. To assess off-target binding, we used a previously described *in vitro* split nanoluc luciferase assay^29^ (**Figure 5A**). Briefly, PYR^NSAID^ is fused to LgBiT and ΔN-HAB1^T+^ is fused to SmBiT. PYR^NSAID^ and ΔN-HAB1^T+^ dimerization increases the local concentration of the two split proteins, reconstituting an active luciferase. Using this assay, we tested 1-100 µM effluent contaminants in addition to a ketoprofen control. While the assay could detect ketoprofen at all concentrations tested, only 100 µM 2,4 dichlorophenol gave a significant luminescence signal over background (**Figure 5B**). As 100 µM is well above typical concentration ranges in effluent, we conclude that PYR^NSAID^ observes minimal background of common effluent chemicals.

**Figure 5.**
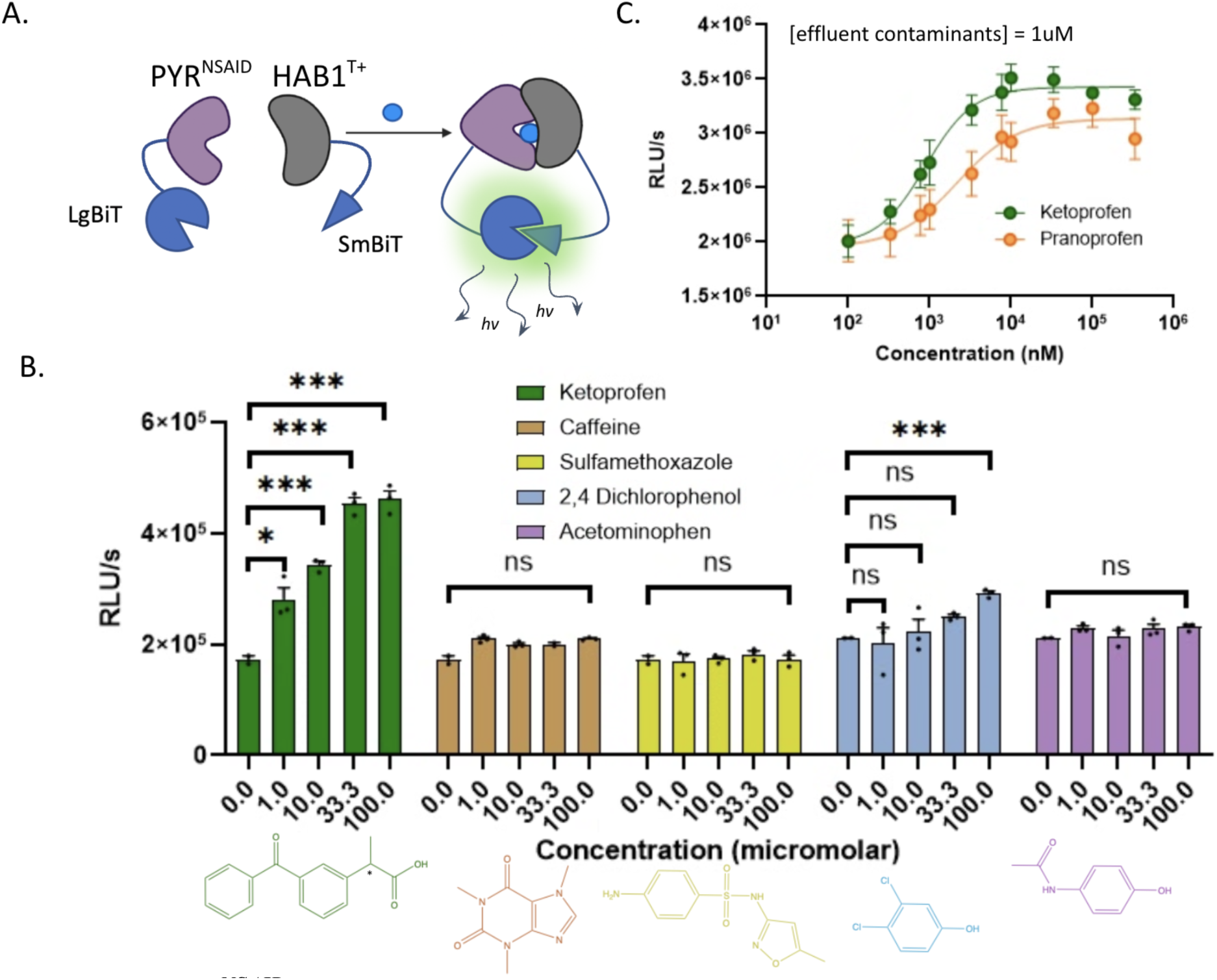
| A PYR1^NSAID^ split luciferase biosensor can sense environmentally relevant NSAIDs with minimal interference by common effluent contaminants. **A.** Schematic of the split luciferase assay. LgBiT and SmBiT are protein fragments from nanoluc luciferase. **B.** RLU/s vs concentration for ketoprofen (green) and other indicated common effluent contaminants. Chemical structures of common wastewater contaminants are shown. For all data, n=3. Error bars represent 1 s.e.m. and in some cases may be smaller than the symbol. ns, not significant; *, p-value < 0.05; ** p-value < 0.01; *** p-value <0.001. **C.** RLU/s vs concentration for ketoprofen (green circles) and pranoprofen (red circles) in the presence of 1 M of common wastewater contaminants (caffeine, sulfamethoxazole, 2,4 dichlorophenol, and acetaminophen).

To determine whether PYR^NSAID^ maintains dose responsive for profens in the presence of these other chemicals, we chose 1µM concentrations of our effluent contaminants to build wastewater standards^46,47^. In ketoprofen and pranoprofen titrations from 100 nM to 330 µM PYR^NSAID^ retained dose responsiveness in these matrices, and the same LODs for ketoprofen and pranoprofen (330 and 770nM, respectively) recorded in the yeast surface display titrations (**Fig 5C**). Thus, PYR^NSAID^ is a profen sensor with minimal activation in the presence of common effluent chemicals and capable of nM-responsive detection of ketoprofen and pranoprofen.

## Discussion

In this study, we used computational and rational protein design, directed evolution, site saturation mutagenesis, and high throughput screening to engineer a protein-based biosensor, PYR^NSAID^, capable of detecting the propionic NSAIDs ketoprofen/dexketoprofen and pranoprofen with nanomolar LODs. PYR^NSAID^ is also sensitive at 33.3 µM and 10 µM to the profens ibuprofen and fenoprofen, respectively. In addition, PYR^NSAID^ exhibits 1 µM sensitivity to diclofenac, an exceedingly toxic acetic acid NSAID^47^ among the most common pharmaceutical pollutants^50^. Finally, we observed a 10 µM limit of detection for tolmetin, another NSAID in the acetic acid class. Using an *in vitro* luminescence assay, PYR^NSAID^ displays minimal activation with other common effluent chemicals and responds to ketoprofen and pranoprofen in dose-responsive manner.

The structural basis by which PYR^NSAID^ can recognize many profens and even acetic acid class NSAIDs is currently unknown. Given that the carboxylate is the only shared functional group between these chemicals capable of hydrogen bonding, we speculate that the likely ketoprofen and pranoprofen alignment involves hydrogen bonds forming between the carboxylic acid in each ligand and the bound water molecule. The proposed alignments of both ketoprofen and pranoprofen in PYR^NSAID^ can be found in **Fig S9**. Whether the protonated carboxylic acid or the carboxylate is involved in binding is unknown. While our yeast and *in vitro* assays were performed at a pH between 7-8, it is well known that carboxylate burial in hydrophobic pockets - like that of the PYR binding pocket - can shift the pKa by several pH units. Regardless, in the absence of a structure, the rationale for why individual mutations found in the deep mutational scan improve profen recognition is hard to rationalize. The most perplexing mutation is V81C given that mutation to the isostere V81S is not beneficial for binding (**Fig 3A-B**). 81C is far removed from other cysteines and deep in the PYR1 binding pocket, and thus unlikely to form a disulfide bond.

The possibilities for a propionic NSAID sensor are considerable; wastewater treatment plants could measure their clearance percentages before releasing into natural waterways. A low cost, rapid, highly selective and sensitive sensor with an output that can be measured using simple UV-Vis would allow water treatment plants to do routine testing at each step of the treatment process, or even continuous monitoring of their workflows to assess NSAID clearance and efficacy of removal techniques. In addition, monitoring of wastewater contaminants serves a purpose in and of itself, as the use of pain management medication reflects fluctuations that occur with changes in seasons, demographics, and water jurisdictions.

There are several steps needed before profen-specific sensors can be applied to environmental monitoring applications. First, PYR^NSAID^ has nM-responsiveness to only ketoprofen/pranoprofen and not all propionic NSAIDs. Further protein engineering can improve the affinity and sensitivity to all environmentally relevant profens. Second, while our sensor exhibited selectivity for a range of propionic and acetic acid NSAIDs, we did observe a significant binding signal in the presence of 100 µM 2,4, dichlorophenol. This concentration is higher than the typical observed range for non-industrial wastewater effluent^48^, in part because triclosan is completely banned in the EU, Canada, Australia, and Japan, and allowed only in toothpaste and hand sanitizer in the US. This limitation, however, may be important to consider for wastewater monitoring of heavily polluted samples like industrial effluent, especially in countries without triclosan bans in effect. Third, we demonstrated sensing using an *in vitro* luciferase and a yeast display assay, each with disadvantages in distributed, real-time sensing applications. Since the PYR-HAB system is a genetically encodable biosensor, alternative *in vivo* and *in vitro* sensing modalities may be better positions for this application^26^. As one example, living cells genetically engineered with luciferase-based PYR-HAB sensing have previously been demonstrated^51^. As the world’s population continues to grow and constraints on our global water supply are compounded, biosensors may play a crucial role in new methods of wastewater monitoring and effluent release into the environment. NSAIDs are an obvious target for wastewater monitoring given their prolific prescription rates, use across multiple industries and organisms, and persistence in the environment. This work provides a genetically encodable sensor for both high quality data collection and quick and reliable efficacy assessment of NSAID removal techniques.

## Acknowledgments

We would like to thank Yves Janin for kindly providing the luciferase prosubstrate, Hikarazine-108. We thank Anika Friedman and Michael Shirts for determining the ketoprofen and pranoprofen conformers. We would also like to thank Nicolle Dunn at the Broomfield, CO wastewater management facility for assisting with effluent standard development and providing useful insight into the wastewater treatment and effluent release process.

## Author contributions

Conceptualization: ZD, SRC, IRW, TAW

Methodology: ZD, HT, TAW Investigation: ZD

Visualization: ZD

Funding acquisition: ZD, SRC, IRW, TAW

Project administration: SRC, IRW, TAW

Supervision: MRS, SRC, IRW, TAW

Writing – original draft: ZD, TAW

Writing – review & editing: ZD, IRW, SRC, TAW

## Data availability

Processed deep sequencing files: **10.5281/zenodo.17782887**

## Code availability

For processing deep sequencing files:

https://github.com/zoedavis7/deep_sequence_reads_processing

