## Supplementary material for "Protein engineering of a genetically encoded biosensor for wastewater detection of profen NSAIDs": Supplmental information

Affiliations:

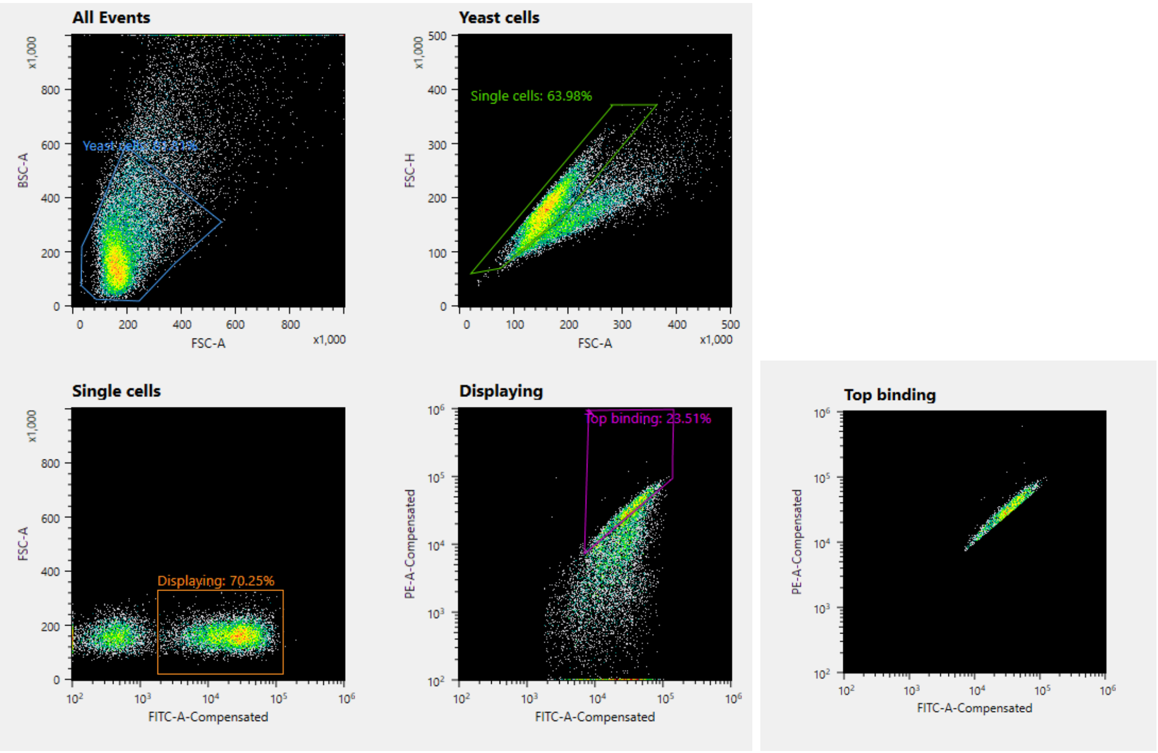


**Figure S1. Example sorting gates used for yeast surface display of the initial propionic acid PYR1 sensor library.** BSC-A (back scatter area) and FSC-A (forward scatter area) are used to gate yeast cells, FSC-H (forward scatter height) and FSC-A are for gating single cells, and FSC-A and FITC fluorescence are used to sort out all displaying cells. “Top binding” is the displaying single yeast cell population with the highest streptavidin-phycoethrin (SAPE) and FITC fluorescence.

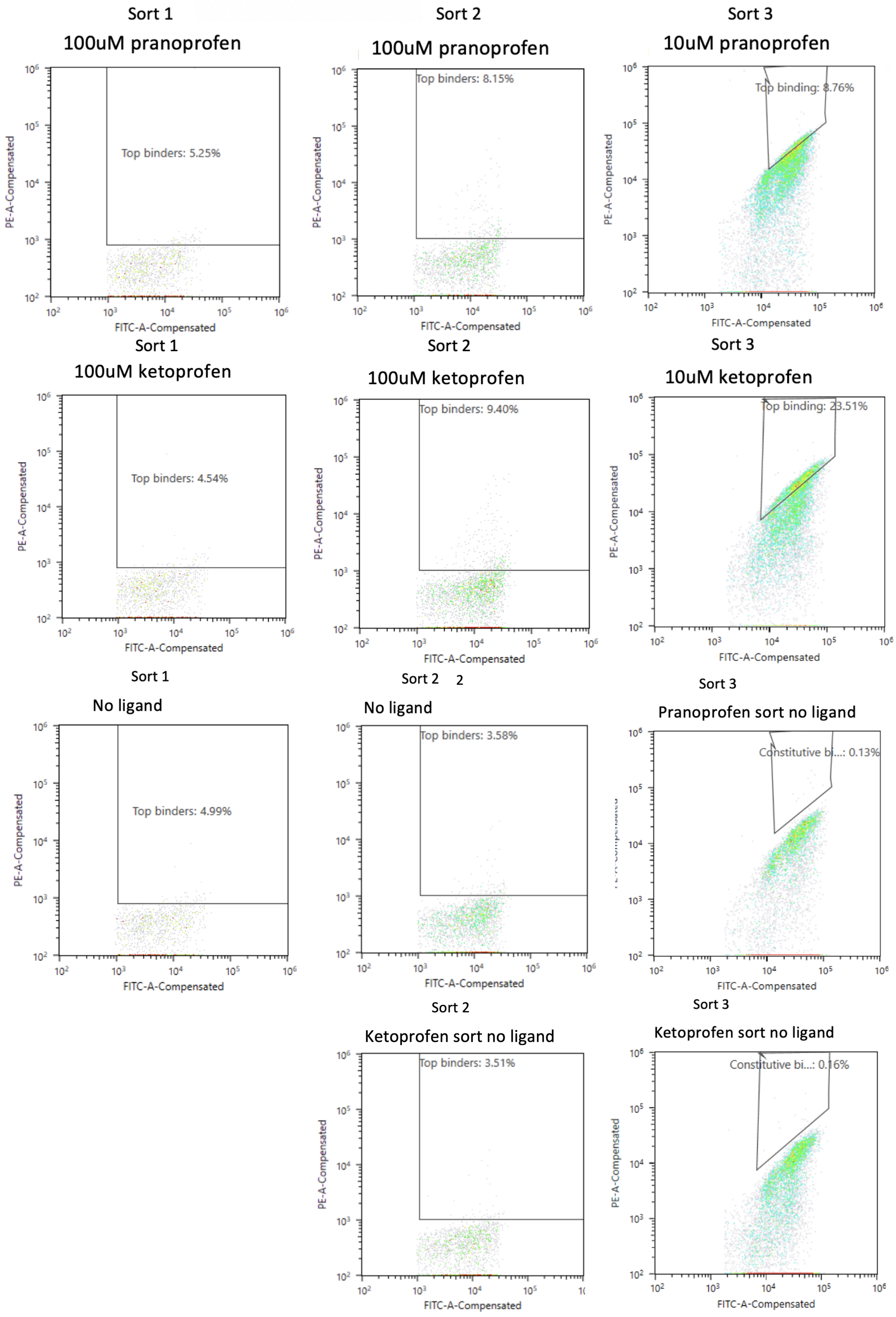


**Figure S2. Cytograms and sorting gates for the first, second, and third rounds of sorting the computationally inspired NSAID library.** Ketoprofen and pranoprofen binders were split into two separate libraries after the first sort. Controls with no ligand (2% (v/v) DMSO) were used to draw gates for each round. Gates uses were FITC-A (fluorescein using a fluorescently conjugated anti-cmyc) and PE-A (using a streptavidin-PE conjugate).

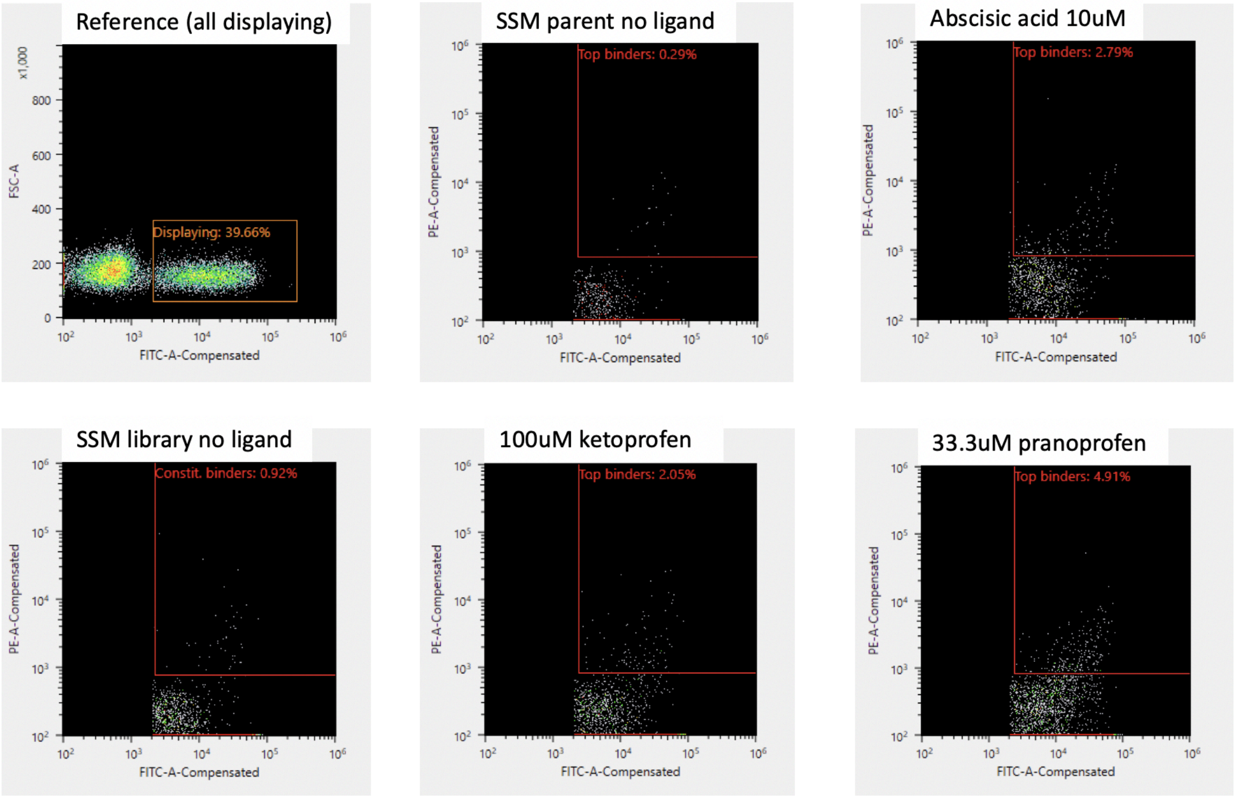


**Figure S3. Cytograms and sorting gates for site saturation mutagenesis library.** 2 (v/v) % DMSO was used for the negative "no ligand" group (Constitutive$^{+}$). No ligand and abscisic acid binding cells were combined in one submission for next generation sequencing, to collectively target all constitutive binding mutations. Biotinylated 300nM ΔN-HAB1$^{T+}$ was used for all binding reactions, followed by secondary labelling with streptavidin-phycoethrin (SAPE). For all data, *n=1*.

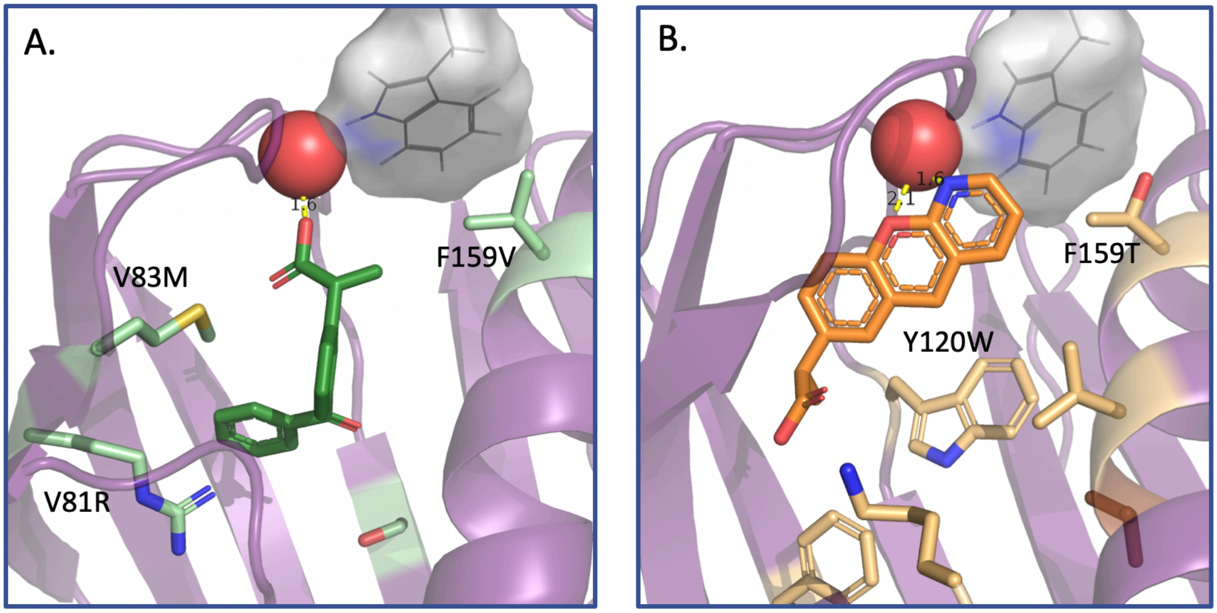


**Figure S4. Two additional alignments in top Y2H hits submitted for a Rosetta design step.** Docked structures of **A.** ketoprofen (forest green, "keto3" hit) and **B.** pranoprofen (orange, "prano5" hit) aligned in the binding pocket of PYR1 (purple). All mutated residues in ketoprofen and pranoprofen hits are colored pale green and gold, respectively. Mutations unique to the top hits shown are identified specifically. Tryptophan 385 of HAB1 is shown in lines with a gray surface. The red sphere at the top of the binding pocket is a bound water essential for the gate-latch-lock chemical induced dimerization mechanism for PYR1-HAB1.

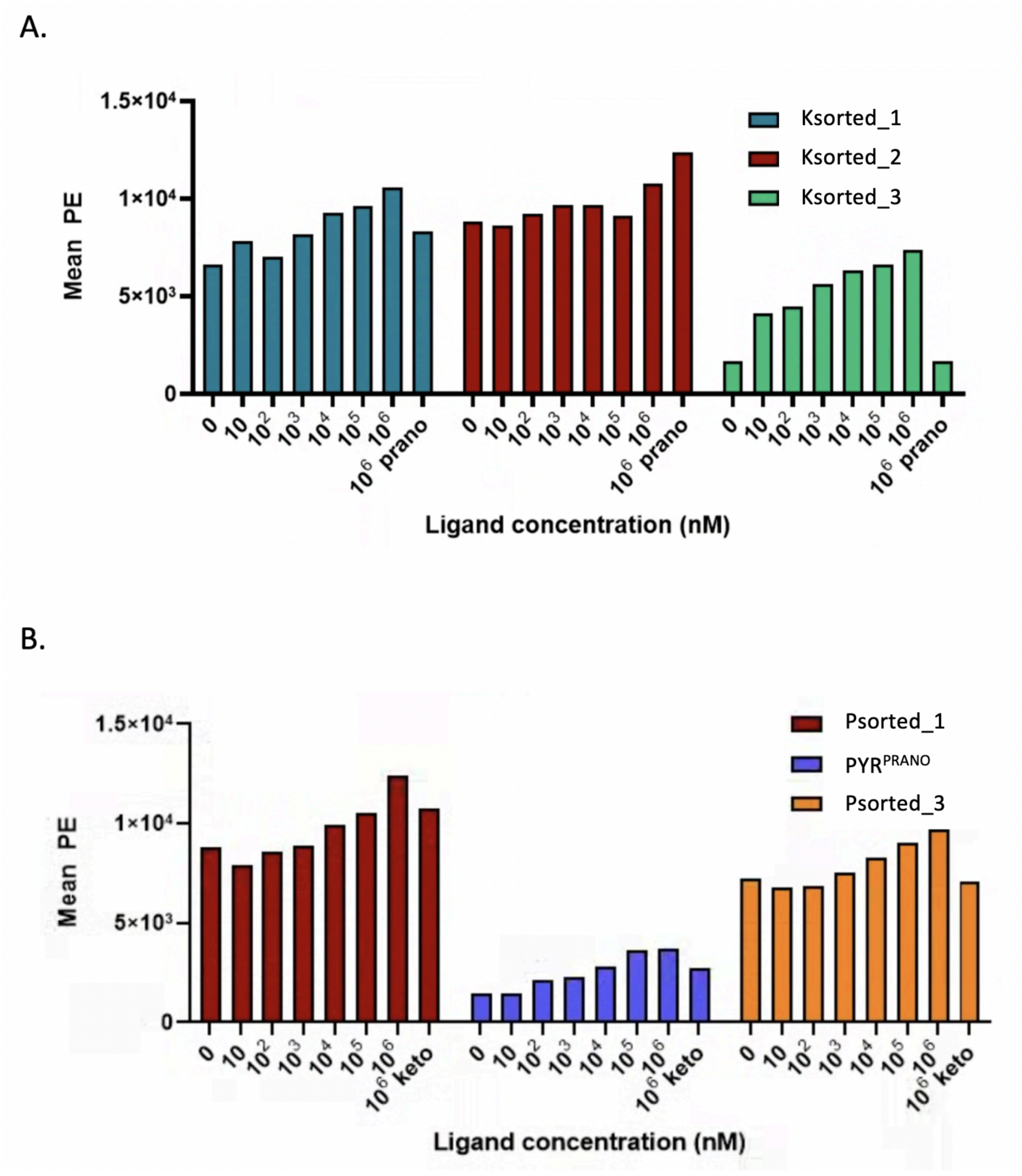


**Figure S5. Yeast surface display binding tests for ketoprofen (A) and pranoprofen (B) hits from the initial NSAID library.** Hits are numbered based on their frequency in next generation sequencing reads. Biotinylated 200nM $\Delta$N-HAB1$^{T+}$ was used for all binding reactions, followed by secondary labelling with streptavidin-phycoethrin (SAPE). For all data, *n=1*.

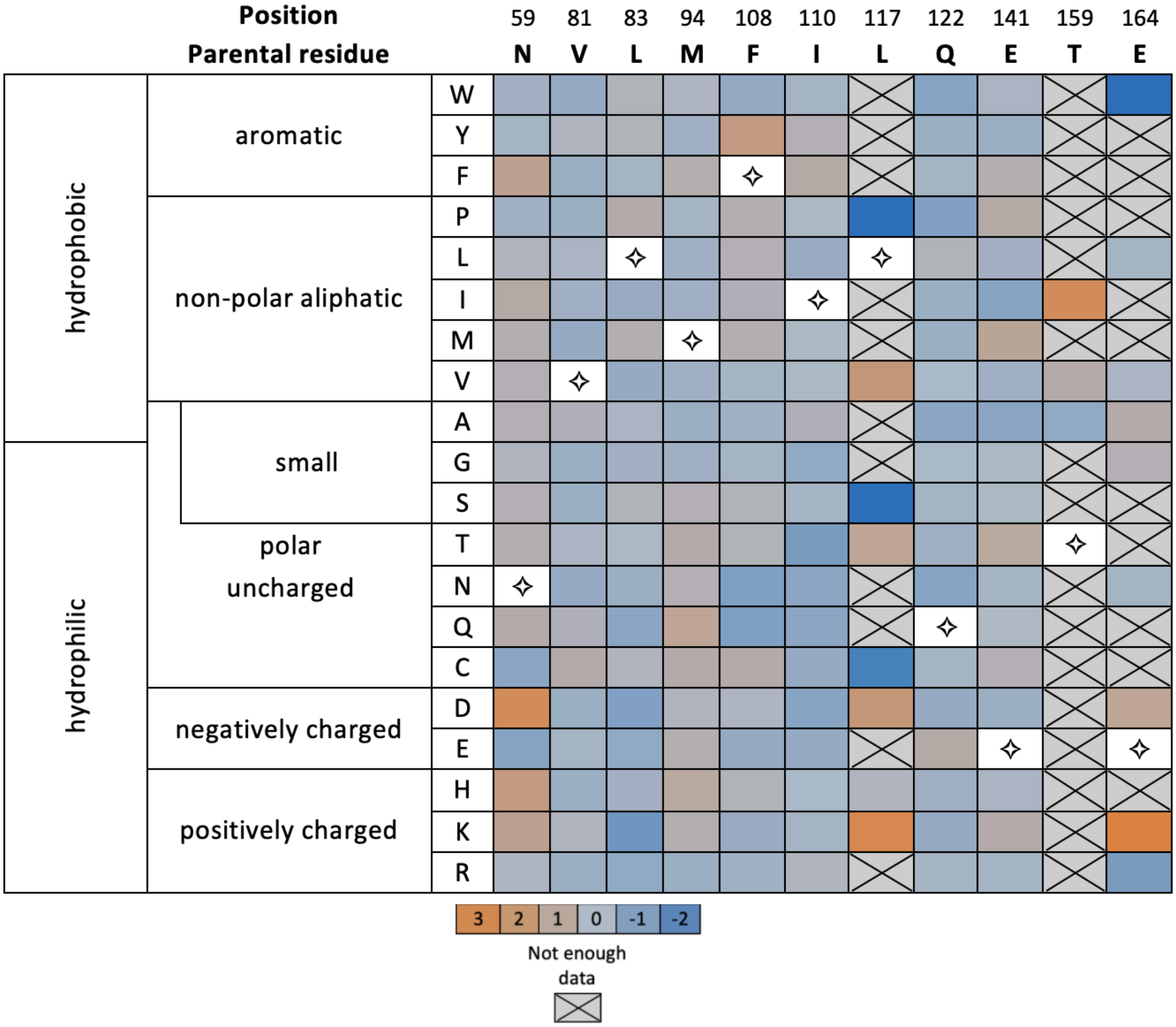


**Figure S6.** Heat map of enrichment ratios for mutations in the combined Constitutive$^{+}$ (2% DMSO) and ABA$^{+}$ (10 $\mu$M abscisic acid) group sorted from the site saturation mutagenesis library.

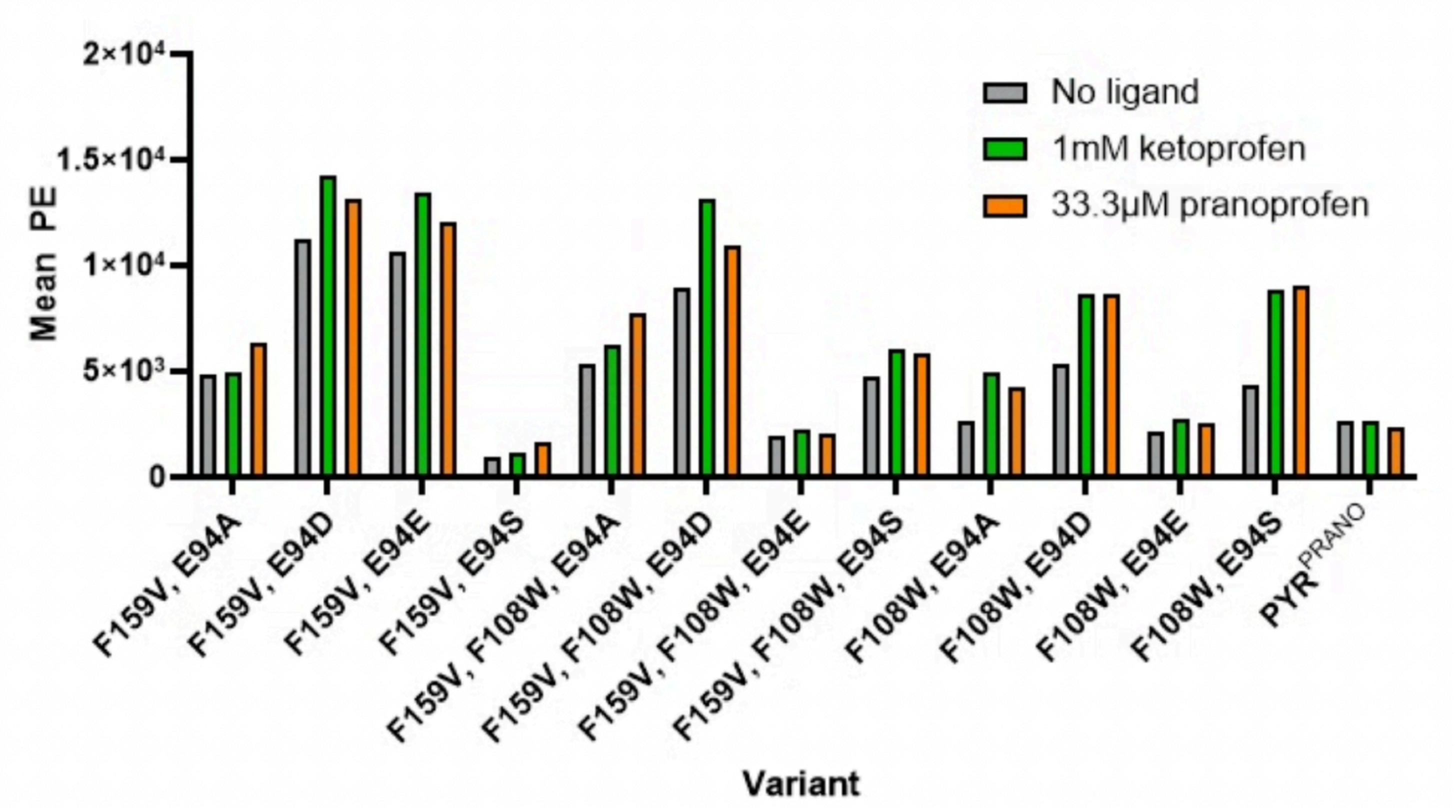


**Figure S7. First round of yeast surface display screening of site saturation mutagenesis hit combinations**. All mutation identities were added in the genetic background of $PYR^{prano}$. Biotinylated 200nM ΔN-$HAB1^{T+}$ was used for all binding reactions, followed by secondary labelling with streptavidin-phycoethrin (SAPE). For all data, *n=1*.

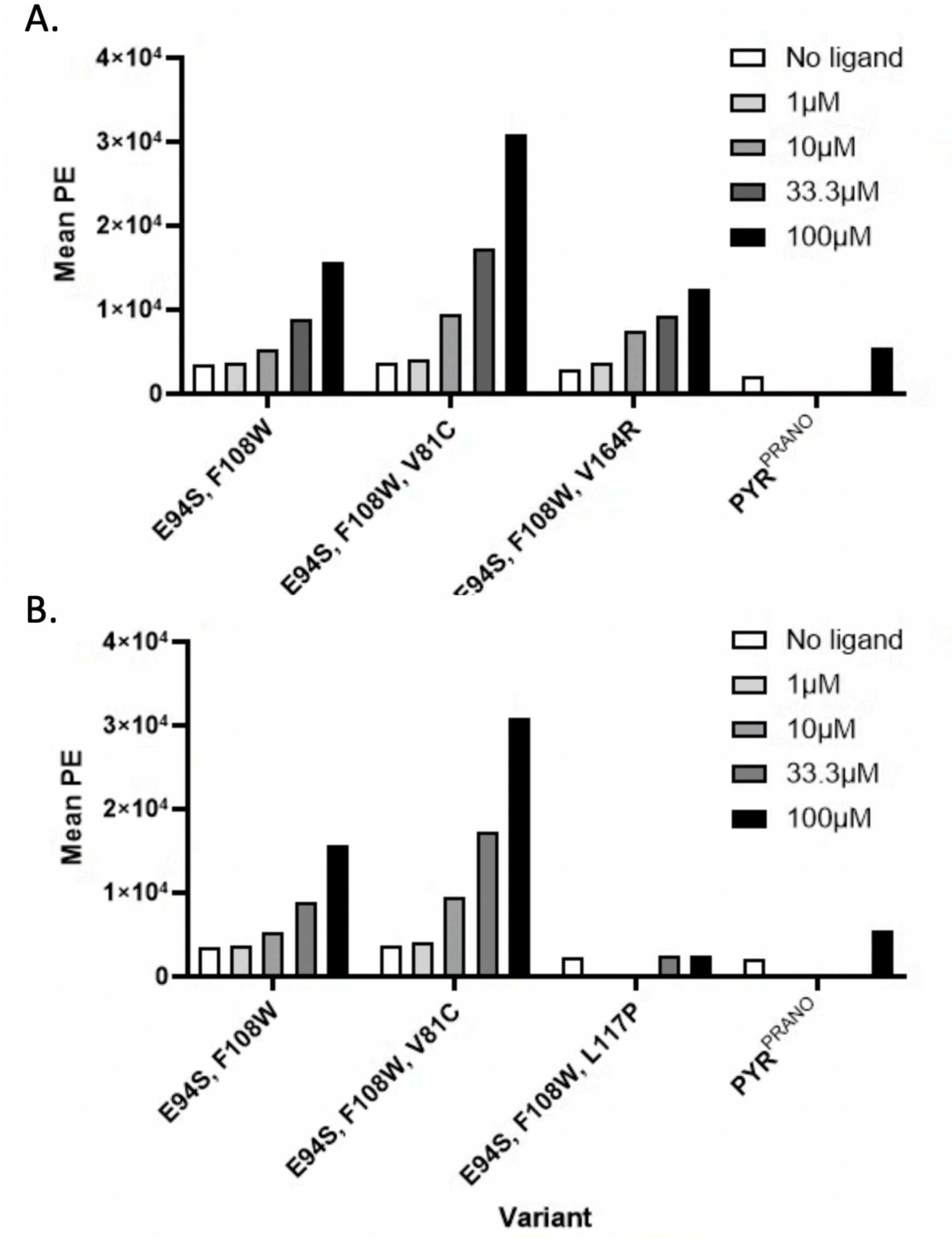


**Figure S8. Second round of yeast surface display screening of site saturation mutagenesis hit combinations.** All mutation identities were added to the existing 4 mutations from the initial library hit, PYR$^{prano}$.This second round of SSM hits did not include the constitutive F159V mutation. Binding to ketoprofen (A) and pranoprofen (B) was tested using biotinylated 200nM ΔN-HAB1$^{T+}$, followed by secondary labelling with streptavidin-phycoethrin (SAPE). For all data, *n=1*.

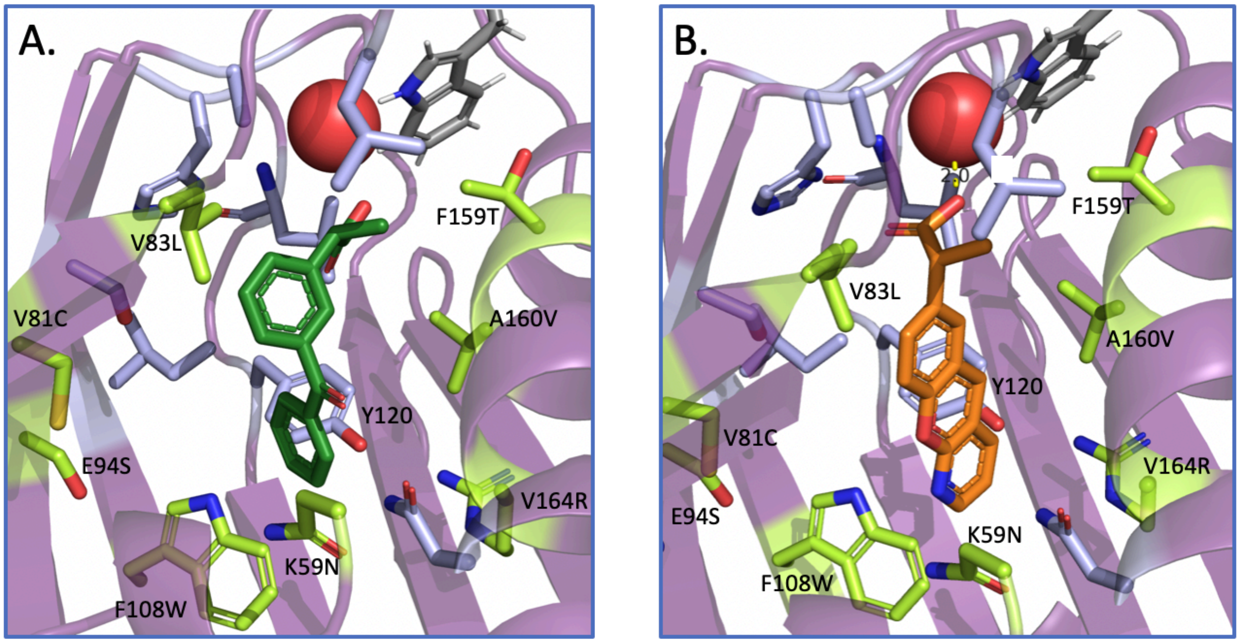


**Figure S9. Proposed alignment of ketoprofen (A) and pranoprofen (B) in our final optimized sensor, PYR$^{NSAID}$.** Docked structures of ketoprofen (forest green) and pranoprofen (orange) aligned in the binding pocket of PYR1 (purple). PYR$^{NSAID}$ mutations relative to the parental background PYR$^{HOT5}$ (K59N, V81C, V83L, E94S, F108W, S122Q, F159T, A160V, V164R) are colored lime and shown as sticks. Other nonmutated binding pocket residues are shown as lavender sticks, and tryptophan 385 of HAB1 is shown as gray sticks. The red sphere at the top of the binding pocket is a bound water essential for the gate-latch-lock chemical induced dimerization mechanism for PYR1-HAB1.

**Table S1.** Complete table of encoded mutations for the first NSAID library.

| Position | 59 | 81 | 83 | 89 | 94 | 108 | 110 | 117 | 120 | 122 | 159 | 160 | 163 | 164 |
|---|---|---|---|---|---|---|---|---|---|---|---|---|---|---|
| $PYR_{WT}$ | K | V | V | A | E | F | I | L | Y | S | F | A | V | V |
| Library | KAIVSND | VR | VRLKMIY | AG | EIVM | FQN | IV | HQN | YW | SQ | VT | AV | VW | VGE |

**Table S2.** Size, transformants and actual coverage of the first NSAID library. % GFP positive is the percentage of GFP positive (no insert) plasmids after Golden Gate assembly; pND003, the destination plasmid used in this work, has a constitutive sfGFP cassette excised during cloning.

| | Initial NSAID library |
|---|---|
| Theoretical library size | 1,354,752 |
| Total # *E. coli* transformants | 590,000 |
| % GFP positive | 17 |
| Total # *S. cerevisiae* transformants | 6,000,000 |
| % Estimated coverage | 30.3 |
| Estimated # of variants | 410,489 |

**Table S3.** Site saturation mutagenesis library coverage for each mutated site in the reference population.

| Position | 59 | 81 | 83 | 92 | 94 | 108 | 110 | 117 | 120 | 122 | 141 | 159 | 160 | 163 | 164 | 167 |
|---|---|---|---|---|---|---|---|---|---|---|---|---|---|---|---|---|
| Observed residues | all | all | all | AL PT | all | all | all | all | *CF HS | all | all | ACIL PRS VW | AEI L | AFI | *ACD GKLN RSVW | DIKS Y |

**Table S4.** Identities of initial Yeast 2 Hybrid hits for pranoprofen and ketoprofen. This information was originally published by Tian et al.[29]

| Variant | Chemical | Position | | | | | | | | | |
|---|---|---|---|---|---|---|---|---|---|---|---|
| | | 59 | 81 | 83 | 108 | 120 | 122 | 159 | 160 | 163 | 164 |
| PYR1 | | **K** | **V** | **V** | **F** | **Y** | **S** | **F** | **A** | **V** | **V** |
| Prano1 | Pranoprofen | K | V | V | **Q** | Y | S | **V** | A | V | V |
| Prano2 | Pranoprofen | K | V | V | F | Y | S | **T** | A | V | V |
| Prano3 | Pranoprofen | **L** | V | **I** | F | Y | S | **V** | A | V | V |
| Prano4 | Pranoprofen | K | V | V | **N** | Y | S | **V** | **V** | V | V |
| Prano5 | Pranoprofen | K | V | V | F | **W** | S | **T** | A | V | V |
| Prano6 | Pranoprofen | K | V | V | F | **W** | S | **T** | A | **W** | **E** |
| Prano7 | Pranoprofen | K | V | V | F | Y | S | **V** | A | V | **G** |
| Keto1 | Ketoprofen | K | **R** | **M** | F | Y | S | **I** | A | V | V |
| Keto2 | Ketoprofen | K | **R** | **L** | F | Y | **Q** | **V** | A | V | V |
| Keto3 | Ketoprofen | K | **R** | **M** | F | Y | S | **V** | A | V | V |

**Table S5.** Top three next generation sequencing hits for each chemical from the initial library.

| Variant | Ligand | % freq. | Position 59 | 81 | 83 | 89 | 94 | 108 | 110 | 117 | 120 | 122 | 159 | 160 | 163 | 164 |
|---|---|---|---|---|---|---|---|---|---|---|---|---|---|---|---|---|
| PYR1 | | -- | K | V | V | A | E | F | I | L | Y | S | F | A | V | V |
| Library | | -- | AIVSND | VR | VRLKMIY | AG | EIVM | FQN | IV | HQN | YW | SQ | VT | AV | VW | VGE |
| Ksorted_1 | keto | 41.9 | K | V | Y | A | E | F | V | L | W | S | V | A | V | V |
| Ksorted_2 | keto | 9.4 | N | V | Y | A | E | F | V | L | Y | S | V | V | V | V |
| Ksorted_3 | keto | 5.3 | S | V | L | A | M | F | V | L | Y | S | V | V | V | V |
| Psorted_1 | prano | 44.7 | N | V | Y | A | E | F | V | L | Y | S | V | V | V | V |
| PYR$^{prano}$ | prano | 21.9 | N | V | L | A | M | F | I | L | Y | Q | T | V | V | E |
| Psorted_3 | prano | 2.7 | N | V | L | G | E | F | I | L | Y | S | V | V | V | V |

**Table S6.** Library size and transformants of the site saturation mutagenesis library.

| | Initial NSAID library |
|---|---|
| Theoretical library size | 320 |
| Total # *E. coli* transformants | 140,000 |
| % GFP positive | 20 |
| Total # *S. cerevisiae* transformants | 1,800,000 |
